# Global skeletal muscle metabolomics reveals mechanisms behind higher response to resistance training in older adults

**DOI:** 10.1101/2025.03.13.642876

**Authors:** Changhyun Lim, Manoel Lixandrão, Dakshat Trivedi, Yun Xu, Konstantinos Prokopidis, Hamilton Roschel, Stuart M. Phillips, Howbeer Muhamadali, Masoud Isanejad

## Abstract

To understand the mechaisnm behind high respond (HighR) compared to low respond (LowR) to resistnace training (RT) and whey protein supplementation (20g/day), we analysied vastus laterails muscle biopsies from a total of 50 participants. Utilising the MRI muscle cross-sectional area (CSA) data, we defined responders as those who had hypertrophy exceeding the 1.7% method error. Quadriceps CSA in the lower responder (LowR) (n=25, mean age 69±5 years) and HighR (n=25, mean age 67±4 years) increased from 53.6 ± 12.1 cm^2^ to 55.4 ± 12.8 cm^2^ after 10 weeks of RET (3.3 ± 1.7%, P < 0.001) and increased the absolute CSA in the higher responders (HighR) from 53.7 ± 12.5 cm^2^ to 59.2 ± 13.6 cm^2^ (10.3 ± 2.0%, P < 0.001). Muscle biopsies were taken from the vastus lateralis before and after RT. We performed untargeted liquid chromatography-mass spectrometry metabolomics to investigate changes in muscle metabolic regulation. The partial least squares discriminant analysis (PLS-DA) yielded the best results using the polar extracts, achieving a 75% average correct classification rate for predicting HighR and LowR. There was no signifncat differences in metabolomic profile at the basline. Our findings revealed several metabolic pathways, including branched-chain amino acid catabolism, tryptophan metabolism (indole and kynurenine pathways), the TCA cycle, gut-derived metabolites, carnitine shuttle metabolism as prominent pathways disrupted in LowR. We provide new insights and has the potential to identify and enhance interventions targeting muscle metabolism, ultimately improving muscle mass and strength to reduce the risk of sarcopenia and frailty in older age.

## 1 INTRODUCTION

Currently, resistance training (RT) is recognized as the most effective ‘anti-frailty’ intervention that can counter muscle loss and improve muscle function in older age [1]. RT involves exercises by a muscle or muscle group against external resistance, typically comprised of higher-load, lower-repetition muscle contractions [2]. RT is known to induce muscle hypertrophy in addition to improving muscular strength and metabolic health. The metabolic adaptations induced by RT result in an increased flux of amino acids and other substrates for the process of protein synthesis and other biosynthetic pathways [3], which are required to increase muscle protein turnover and mass [1, 4]. Despite the existing evidence suggesting a broad metabolic change with RT[3, 5], less is known about the effect of RT at the muscle metabolic level, especially in aging muscle. Biochemical assessments, including metabolomics in response to exercise, have the potential to identify biological pathways that are essential in developing strategies for the management of ageing diseases and frailty [6, 7]. We conducted high-throughput untargeted liquid chromatography-mass spectrometry on vastus lateralis muscle biopsies for metabolic profiling. These biopsies were taken 10 weeks apart from a well-phenotyped older population that had completed a resistance training (RT) intervention [8]. From this analysis, we identified distinct phenotypes: lower responders (LowR; bottom tertile) and higher responders (HighR; top tertile) based on hypertrophy profiles following the RT intervention. We aimed to relate muscle metabolic phenotypes before and after 10wk of RT and, based on the HighR compared to LowR phenotype, to assess whether associated biochemical networks related to biological response associated with the response to the application of the loading stimulus, which is well-known to improve muscle function, counter anabolic resistance and enhance muscle function.

## 2 METHODS

### 2.1 Study population

Of the 83 participants in the previous study [8], muscle biopsies were obtained from 67 of the healthy older adults, comprising 30 men (68 ± 4 years, BMI = 26.4 ± 3.0 kg/m^2^) and 37 women (68 ± 5 years old, BMI = 26.4 ± 4.3 kg/m^2^). The participants were generally healthy, but some had metabolic diseases (type 2 diabetes, hypertension, hypercholesterolemia) for which they were receiving medications. None had regularly engaged in structured exercise, such as resistance or aerobic training, for at least six months prior to the study. Exclusion criteria included type I diabetes, ischemic myocardial disease, arrhythmia, uncontrolled hypertension, and any major orthopedic or musculoskeletal disorder. The study was approved by the Human Research Ethics Committee of the University of Sao Paulo. All participants provided written, informed content prior to taking part in the study and to have their tissue samples stored and analyzed after the main study. The main trial was registered at clinical trial registration number NCT06718712. The ethical approval for muscle biopsy analysis was obtained from the University of Liverpool Central University Research Ethics Committee, reference number 12689.

### 2.2 Resistance exercise training protocol

The protocol for RT and dietary assessments has been described previously [8]. In summary, all participants performed unilateral knee extension exercises two times a week for 10 weeks. One of the subject’s legs performed 1 set, while the contralateral leg performed 4 sets of 8-15 repetitions, with 90 seconds of rest interval between sets. In the present study, we focused exclusively on the leg that performed 4 sets (i.e., the multiset condition), which was previously shown to show a more robust muscle hypertrophy response [8].

### 2.3 Hypertrophy response classification

To classify response variation, we used the change in MRI-measured muscle cross-sectional area (CSA) from baseline and compared that value to the usual method variation in magnetic resonance imaging (MRI) measurement method error. The method error on repeated scans was 1.7%. When we rank-ordered participants’ hypertrophy for males and females separately from baseline, the 67 participants from whom we had muscle were divided, and two distinct groups were identified comprising the top 25 (n=10 males and n=13 females) and bottom 25 (n=15 males and n=12 females) participants in terms of their rank from the lowest responses for the males and females (Figures 1A, B). We only used the hypertrophic response to 4 sets of exercise as it resulted in a more robust hypertrophic response, but responder status was conserved in the same participants within the leg that performed only one set, but admittedly with a less robust anabolic response (data not shown).

**Figure 1.**
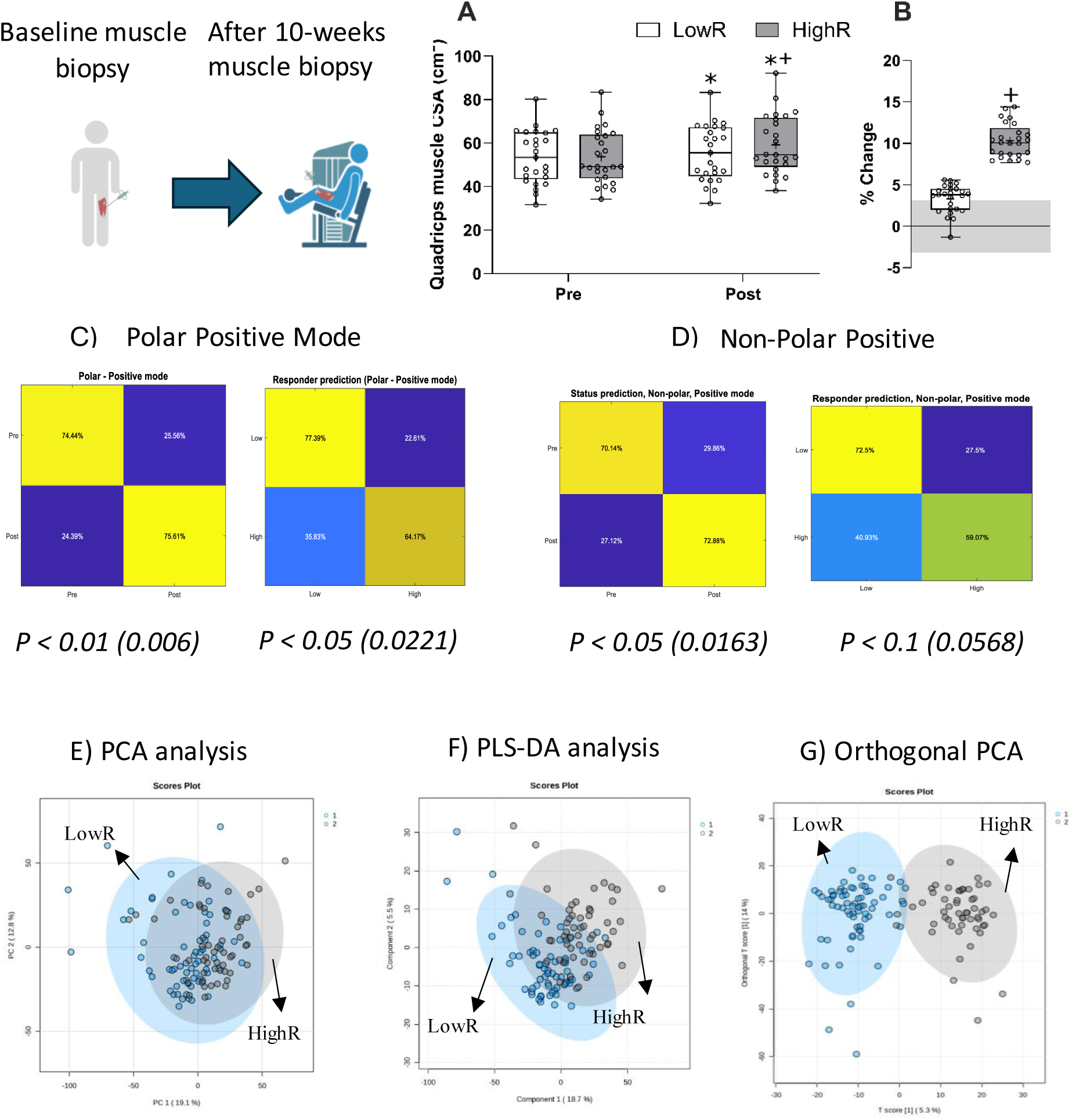
Study design overview and analytical methods. A) Muscle biopsies were taken from participants before and after 10 weeks of Resistance Training, magnetic resonance imaging-measured quadriceps muscle cross-sectional area (CSA), from which the high responders (HighR) and low responders (LowR) groups were established. B) The change in the quadriceps muscle CSA between HighR and LowR, * significantly different from pre-training; + significantly different from LowR (P<0.001). The shaded box shows normal method variation. Panels C) and D) Show the polar positive and non-polar positive PLS-DA analysis. MetaboAnalyst version 6.0 was used to visualise E) Principal components analysis (PCA), F) Partial Least Squares Discriminant Analysis (PLSDA) and G) Orthogonal PCA.

### 2.4 Untargeted liquid chromatography-mass spectrometry (LCMS) metabolomics analysis

#### 2.4.1 Sample preparation

Reagents used included Optima^®^ LC-MS grade methanol (MeOH), chloroform (CHCl_3_), acetonitrile (ACN), isopropyl alcohol (IPA) and water (H_2_O) were used for sample preparation and subsequent analysis. LC-MS LiChropur™ grade (≥ 99.0%) ammonium formate and formic acid (FA) were used as solvent buffers. HPLC grade absolute ethanol (99.8% v/v) was used for maintenance and cleaning. Briefly, 10 mg (±1 mg) of muscle tissue was suspended in 216 μL of a pre-chilled MeOH: Water (2.86:1) mix in 2 mL Precellys^™^ tubes. Samples were homogenised in two consecutive 10-s cycles at 5000 rpm using a Qiagen PowerLyzer^®^ 24 (Qiagen, Germany). The homogenised samples were transferred to 2 mL Eppendorf tubes^®^ and mixed with 240 μL of pre-chilled CHCl_3_: H_2_O (2:1) and allowed to incubate on ice for 10 min. Samples were vortexed (30 s) and centrifuged (3500 *g*, 4 °C, 5 min) to achieve phase separation. Aliquots (200 μL) from each of the polar (upper) and non-polar (lower) phases were collected in separate 2 mL Eppendorf tubes^®^ and dried to evaporation in a cold-trap vacuum centrifuge. The dried extracts were stored at -80 °C until UHPLC-MS analysis. Metabolomic analysis was carried out th the Cetnre of metabolomic Research, Liverpool Shared Research Facilty (LIV-SRF), the Univertity of Liverpool.

#### 2.4.2 Quality control and assurance (QA/QC)

A fixed amount of homogenized tissue extract was aliquoted from all samples to prepare pooled QC samples and conditioning QC samples as described in [9]. Pooled QCs were used for the assessment of data quality. Extraction blanks, solvent blanks, QC samples and system suitability test (SST) samples were prepared alongside samples using the same preparation method [9].

#### 2.4.3 UHPLC-MS analysis

Untargeted metabolomics data were acquired using a ThermoFisher Scientific Vanquish UHPLC system coupled to an Orbitrap ID-X Tribid mass spectrometer (MS) (ThermoFisher Scientific Inc., UK). Dried polar extracts were reconstituted in 100 μL of ACN: Water (90:10), and the non-polar extracts were reconstituted in 100 μL of Water: Methanol (80:20) and transferred to glass vials. The polar extracts were separated on a Hypersil GOLD™ aQ (C_18_, polar 2.1 mm x 10 mm x 1.9 μm, ThermoFisher Scientific) while the non-polar extracts were separated on Hypersil GOLD™ Vanquish ™ C_18_ column. During analysis, columns were maintained at 50 °Cover a 15 min gradient elution (detailed in Table (X)) and 0.4 mL/min flow rate. Samples were stored at 4 °C throughout the run in LC-autosampler. A 5 μL sample was injected for both electrospray ionisation modes (ESI+/-). Data were acquired in full-scan mode. Each analytical batch was bracketed with blank and conditioning QC injections and contained periodic pooled QC sample injections[10]. To aid compound annotation and identification ddMS^2^ data were acquired from 3x QC samples over (a) 66.7-1000, (b) 66.7-300, (c) 300-600 and (d) 600-900 *m/z* ranges[11]. Detailed gradient elution profiles, solvent composition (Supplementary Table 1) and MS parameters are described in (Supplementary Table 2) for all modes.

### 2.5 Statistical analysis

The distribution of data was assessed using the Shapiro-Wilk test. Baseline characteristics of the participants among higher and lower responder groups were analyzed using an unpaired T-test. A Chi-square test was performed for nominal variables (i.e. clinical conditions in the present study). Changes in absolute quadriceps CSA between groups were assessed using a linear mixed model (group x time), with a Tukey post hoc test performed when a significant interaction was detected. The percent change in the CSA between higher and lower groups was analyzed using an unpaired t-test. Statistical significance was set at P < 0.05, and data are presented as mean ± SD. Statistical analyses were completed using R (version 4.3.2).

### 2.6 Metabolomic analysis

The LC-MS data were first deconvolved using compound discoverer 3.2 (Thermo-Fisher). Tentative annotations were made for each detected metabolic feature using MS/MS spectra matching. The deconvolved data matrices were then imported into MATLAB (Mathworks, MA) 2023a for multivariate analysis. Principal component analysis was performed to inspect the integrity of the data. Partial least squares-discriminant analysis (PLS-DA) model was then employed to discriminate high responder vs. low responder and also pre-treatment vs. post-treatment. The models were validated by using 1,000 bootstrapping procedures as described previously [12]. To assess the statistical significance level of the performance of the models, permutation tests were also performed. Further, we manually curated all metabolites to discern biologically relevant signals from noise. Lastly, N-way ANOVA was performed on each log-transformed metabolic feature to detect whether there are statistically significant differences between pre-treatment vs. post-treatment or high responder vs. low responders. A Benjamini-Hochberg false discovery rate was then employed on the p-values of the ANOVA tests to control the increased risk of false discovery by multi-testing. Further, to visualize the results of PCA analyses, MetaboAnlyst 6.0 was used to present the PCA, PLSDA and orthogonal PCA scores plots (Figure 1E, F, G). Orthogonal PCA has a similar predictive capacity compared to PLS and improves the interpretation of the predictive components and the systematic variation [13].

## 3 RESULTS

Participants’ characteristics for LowR and HighR are shown in Table 1. Of the total participants (n=50), 25 were classified as lower responders (LowR) and 25 as higher responders (HighR,), based on a 2% (normal method variability) cut-off value (Figure 1). No significant differences were observed in anthropometric measures between the groups prior to the intervention (P > 0.05; Table 1). Additionally, no significant differences were found in the prevalence of clinical conditions (T2DM, Hypertension, and cholesterol) or habitual nutritional intake (data not shown) across the groups (P > 0.05). The absolute quadriceps CSA in the LowR increased from 53.6 ± 12.1 cm^2^ to 55.4 ± 12.8 cm^2^ after 10 weeks of RET (3.3 ± 1.7%, P < 0.01; Figure 1A). The 10 weeks of RET also increased the absolute CSA in the HighR from 53.7 ± 12.5 cm^2^ to 59.2 ± 13.6 cm^2^ (10.3 ± 2.0%, P < 0.001; Figure 1B). While both LowR and HighR showed increases in the CSA following the RET, the percent change in CSA was statistically significantly, and arguably physiologically, greater in the HighR compared to the LowR (P < 0.001; Figure 1B).

**Table 1.**
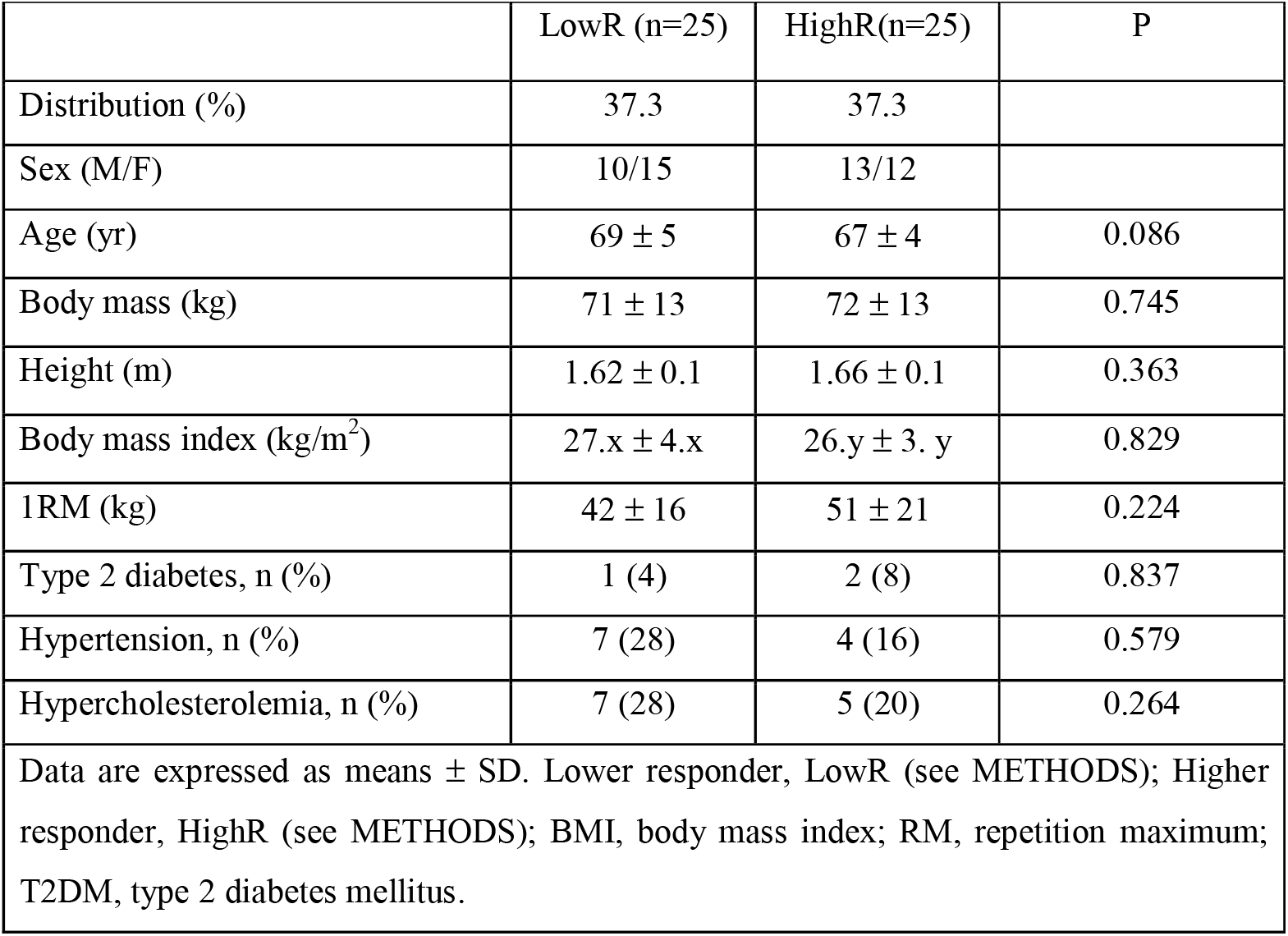
Baseline characteristics of the participants.

| **Table 1. Baseline characteristics of the participants.** |  |  |  |
| --- | --- | --- | --- |
|  | LowR (n=25) | HighR(n=25) | P |
| Distribution (%) | 37.3 | 37.3 |  |
| Sex (M/F) | 10/15 | 13/12 |  |
| Age (yr) | 69 ± 5 | 67 ± 4 | 0.086 |
| Body mass (kg) | 71 ± 13 | 72 ± 13 | 0.745 |
| Height (m) | 1.62 ± 0.1 | 1.66 ± 0.1 | 0.363 |
| Body mass index (kg/m<sup>2</sup>) | 27.x ± 4.x | 26.y ± 3. y | 0.829 |
| 1RM (kg) | 42 ± 16 | 51 ± 21 | 0.224 |
| Type 2 diabetes, n (%) | 1 (4) | 2 (8) | 0.837 |
| Hypertension, n (%) | 7 (28) | 4 (16) | 0.579 |
| Hypercholesterolemia, n (%) | 7 (28) | 5 (20) | 0.264 |
| Data are expressed as means ± SD. Lower responder, LowR (see METHODS); Higher responder, HighR (see METHODS); BMI, body mass index; RM, repetition maximum; T2DM, type 2 diabetes mellitus. |  |  |  |

### 3.1 Analysis of muscle metabolome in response to RT

At study entry (baseline), there were no significant differences in muscle metabolome between groups. From the PLSDA analysis best results were obtained from the models using polar extracts in predicting HighR and LowR response (Figure 1C-G) with 75% averaged correct classification rate (CCR) in predicting pre- or post-RT. The corresponding empirical p-value of 0.006 (i.e., only 6 out of 1,000 permutation tests had resulted in the NULL models obtaining better results than that of the corresponding observed models). The same averaged CCR of models using non-polar extracts was slightly lower, at 72%, with an empirical p-value of 0.0163, which is still statistically significant (Figure 1D). The performances of the models predicting LowR vs. HighR were generally worse than those predicting Pre vs. Post. The VIP included mostly such as amino acids, peptides include BCAAs, typtophan, proline, and glutamine. From the PLSDA results, we also developed the variable importance in projection (VIP) as presented in Figures 2 A and B. Further, the top enriched pathways are shown in Figures 2C and 2D (the data will be deposited MetaboLights).The averaged CCR in predicting LowR vs. HighR of the models using polar metabolites (Figure 2B) was 71% (p = 0.0221), and that of the models using non-polar metabolites (Figure 2B) was 66% (p = 0.0568). We observed distinct metabolic insights from the analysis of HighR compared to LowR. In this study, we have provided an overview of our findings based on the most prominent functional features of these metabolites. In particular, the HighR metabolic phenotype showed greater relative levels of amino acids (isoleucine, leucine, valine, phenylalanine, lysine, glutamine, methionine, tyrosine, and citrulline) (Figure 3A and B); peptides and other related metabolites (such as carnosine, acetylcarnitine, creatine, alanyl-glutamine, taurine, and spermidine; Figure 3C); vitamins and cofactors such as riboflavin and nicotinamide (Figure 4A); gut-related metabolites (indole, kynurenic acid, adrenaline, and isoprenaline; Figure 4B); neurotransmitters and signalling molecules (such as acetylcholine and adenosine diphosphate); and finally, fatty acid derivatives such as alpha-aminoadipic acid and isoprene (Figure 4C).

**Figure 2.**
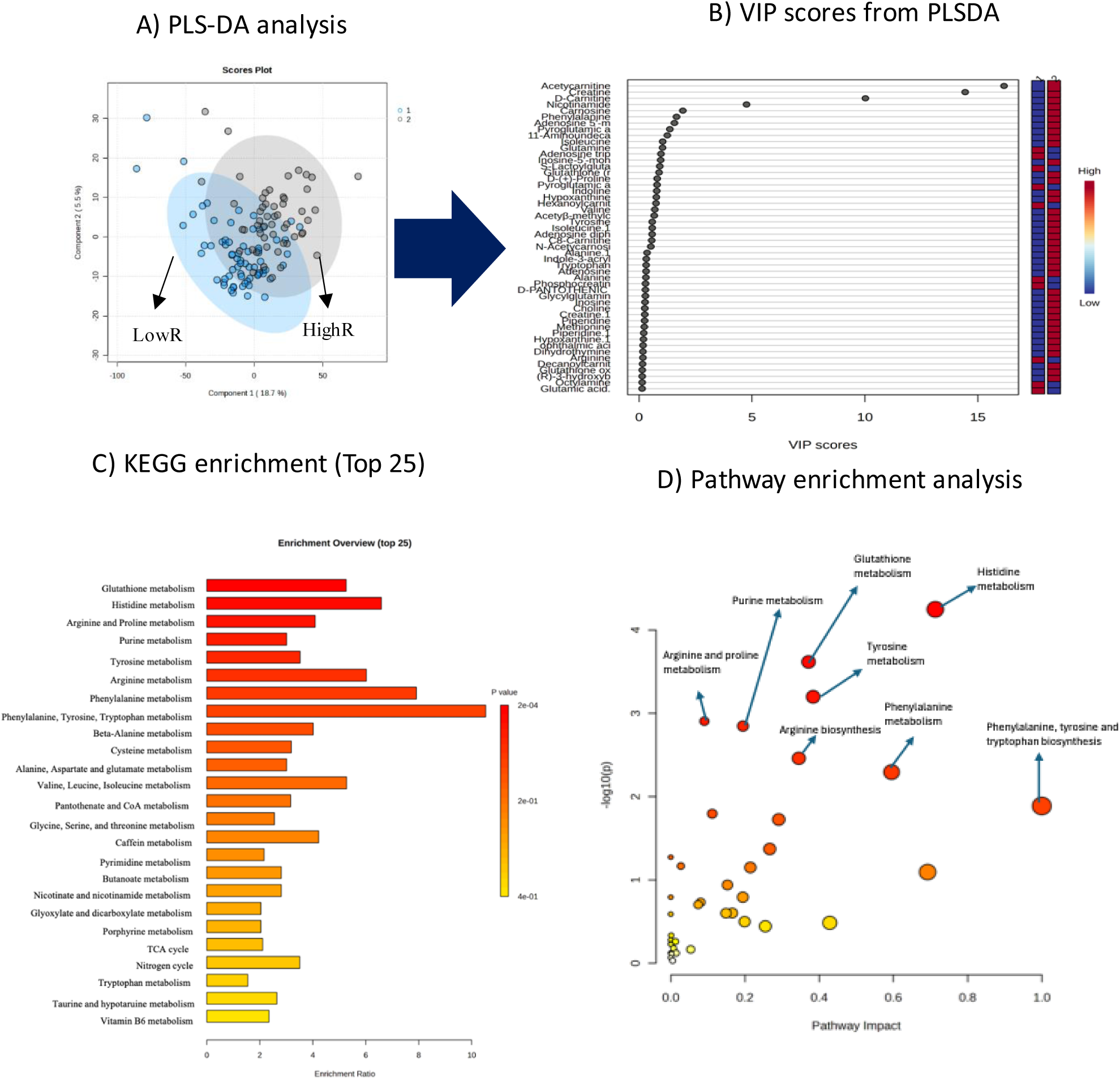
Study design and analytical methods. A) Supervised Partial Least-Squares Discriminant Analysis (PLS-DA) was also used to calculate the top 50 metabolites. B) Variable importance in projection. C) Shows the results of pathways enrichment analysis by using the name of the compound identified from the KEGG database. D) Shows the enrichment analysis impact and p-values indicating the top enriched pathways. All analyses were performed utilising MetaboAnalyst version 6.0, according to 1250 sub-chemical class metabolite sets of library-matched RaMP-DB (integrating KEGG via HMDB, Reactome, WikiPathways).

**Figure 3.**
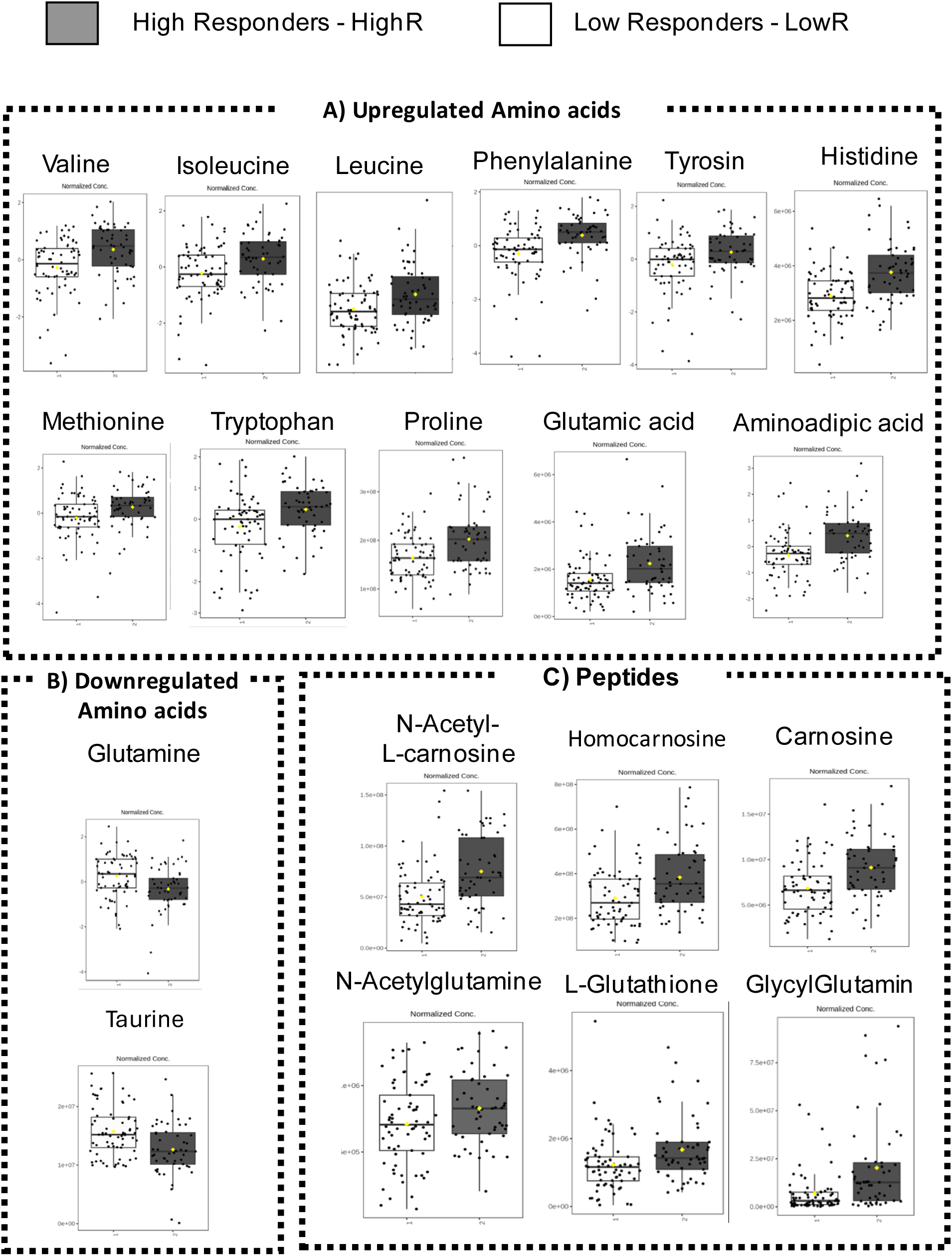
Upregulated and downregulated amino acids and peptides. (given the large number of metabolites, these box plots represent a selected subset). Multiple t-tests were performed in MetaboAnalyst 6.0. The box plots show A) upregulated Amino Acids, B) Downregulated amino acids, and C) Upregulated Peptides with FDR<0.05 that were upregulated or downregulated, comparing HighR to LowR. Given the large number of metabolites these box plots are representing a selected number

**Figure 4.**
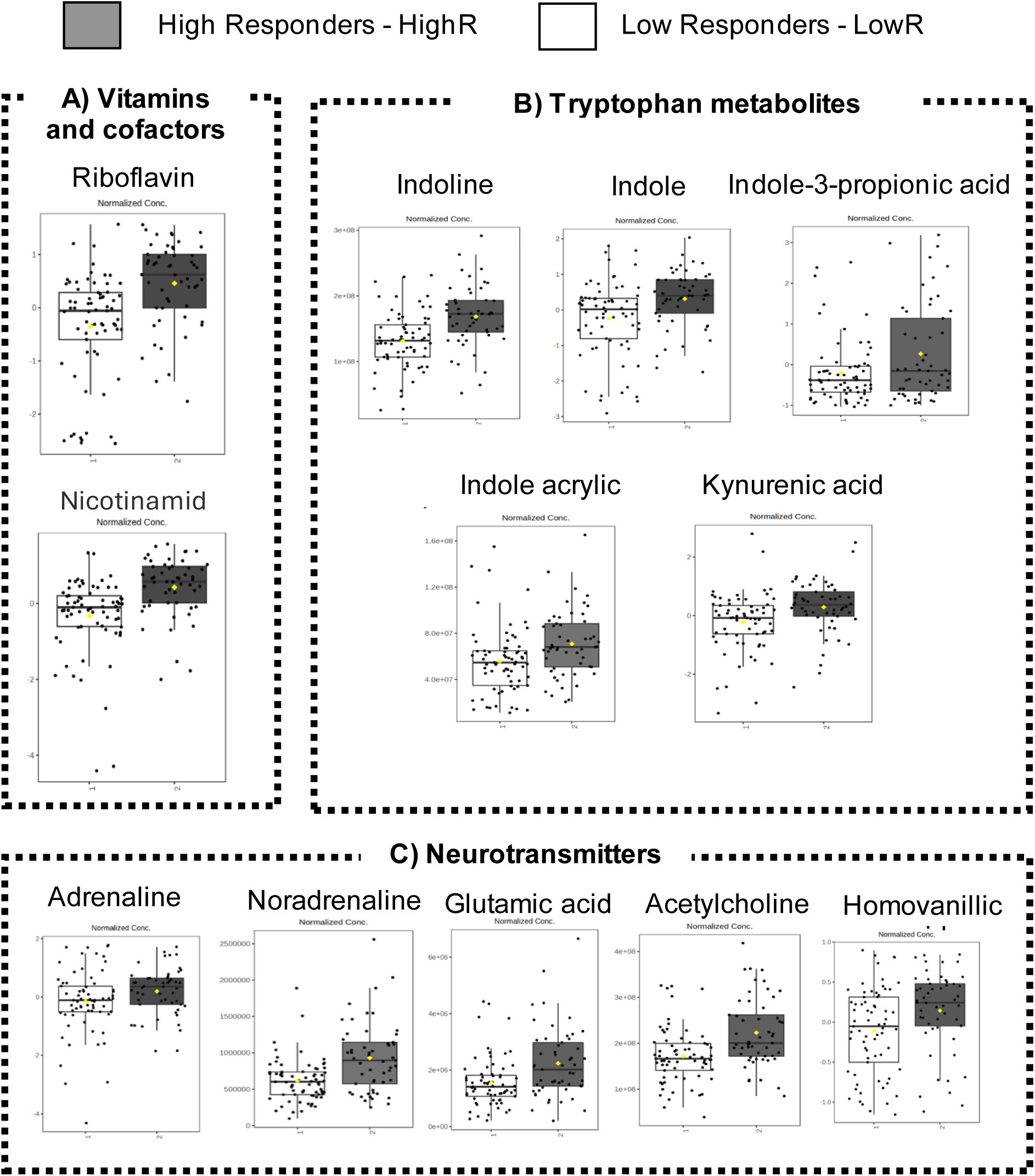
Upregulated and downregulated vitamins, Tryptophan-related gut metabolites, and neurotransmitters. (given the large number of metabolites, these box plots represent a selected subset). Multiple t-tests were performed in MetaboAnalyst 6.0. The box plots show examples of significant metabolites between high responders compared to low responders, grouped based on their function. A) Vitamins and co-factors, B) Gut-related metabolites, and C) Neurotransmitters. These metabolites are either from VIP or with biological relevance with FDR<0.05 that were upregulated, comparing HighR to LowR.

### 3.2 Pathway analysis

The pathway analysis indicated that greater impact can be attributed to peptides and amino acid metabolism. (Figure 2C, D). The topmost significant pathways, as indicated by the lowest p-values, include histidine, glutathione, tryptophan and tyrosine metabolisms (Figure 2C). Further, we utilized the KEGG pathways and visualised branched-chain amino acids (BCAA) pathways (Figure 5) and tryptophan metabolism in the muscles (Figure 6) and their response to long-term RT. We identified valine, leucine isoleucine and their branched-chain aminotransferase metabolites, and branched-chain fatty acids being upregulated in response to exercise (Figure 5). Furthermore, the results suggested that both tryptophan-kynurenine and tryptophan-indole metabolism were significantly upregulated in response to RT in human muscle (Figure 6). Results from the functional Mummichog pathway analysis are presented in Figure 7, and full data is available in Supplementary material 1 and visualised in Figure 7 A. The top 5 most enriched Mummichog pathways in HighR compared to LowR included tyrosine, aspartate and tryptophan metabolisms (Figure 7B).

**Figure 5.**
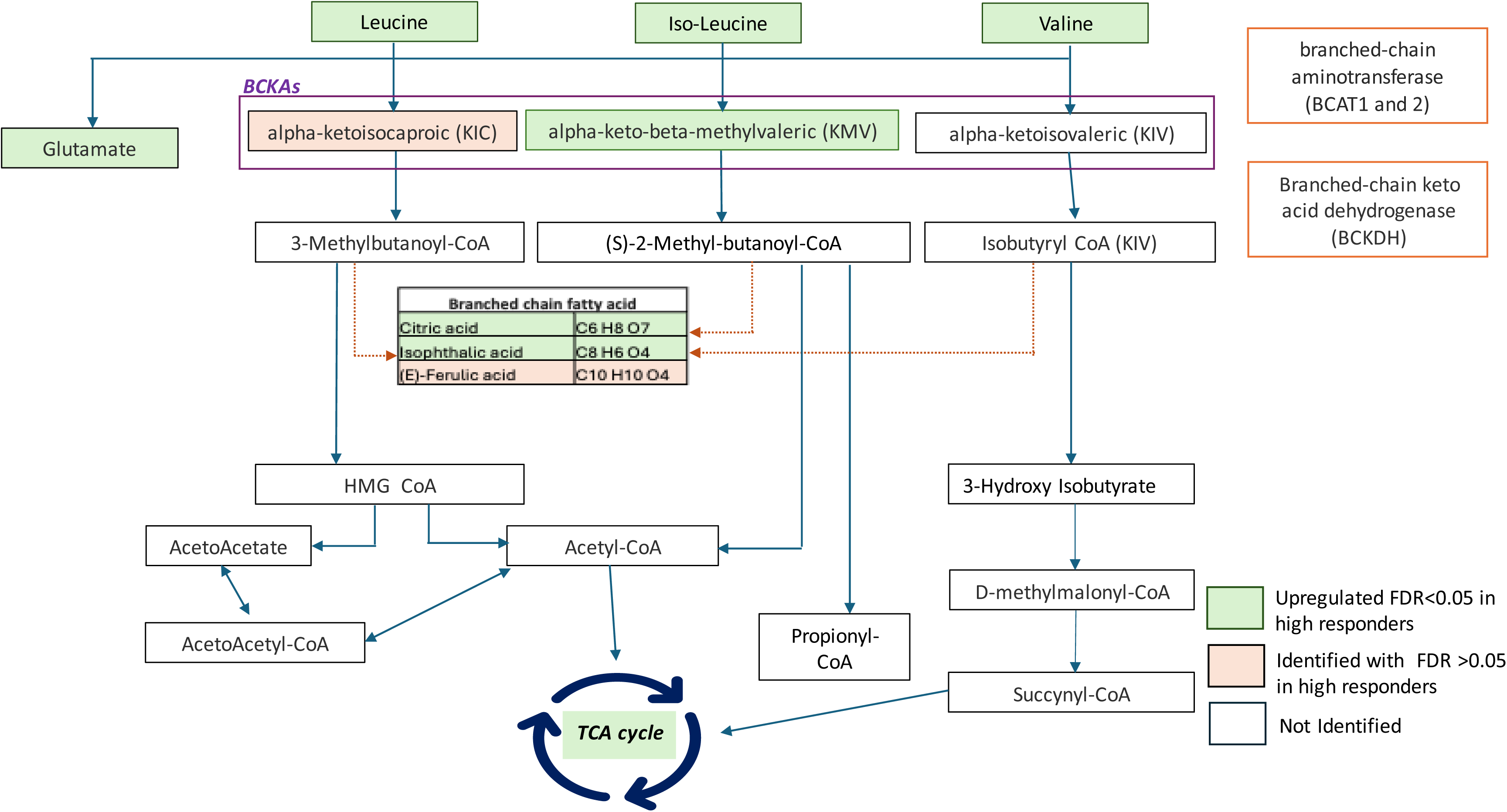
Schematic overview of the Branc Chain Amino Acids-related pathways and the significant metabolites detected in this study adapeted from KEGG. Illustration of the catabolism pathways of BCAAs (leucine, isoleucine, and valine). The significant metabolites identified in this study were overlaid onto the BCCA pathway, to assess its components based on their response to resistance training. The false discovery rates (FDRs) were derived from multiple t-tests conducted in MetaboAnalyst 6.0.

**Figure 6.**
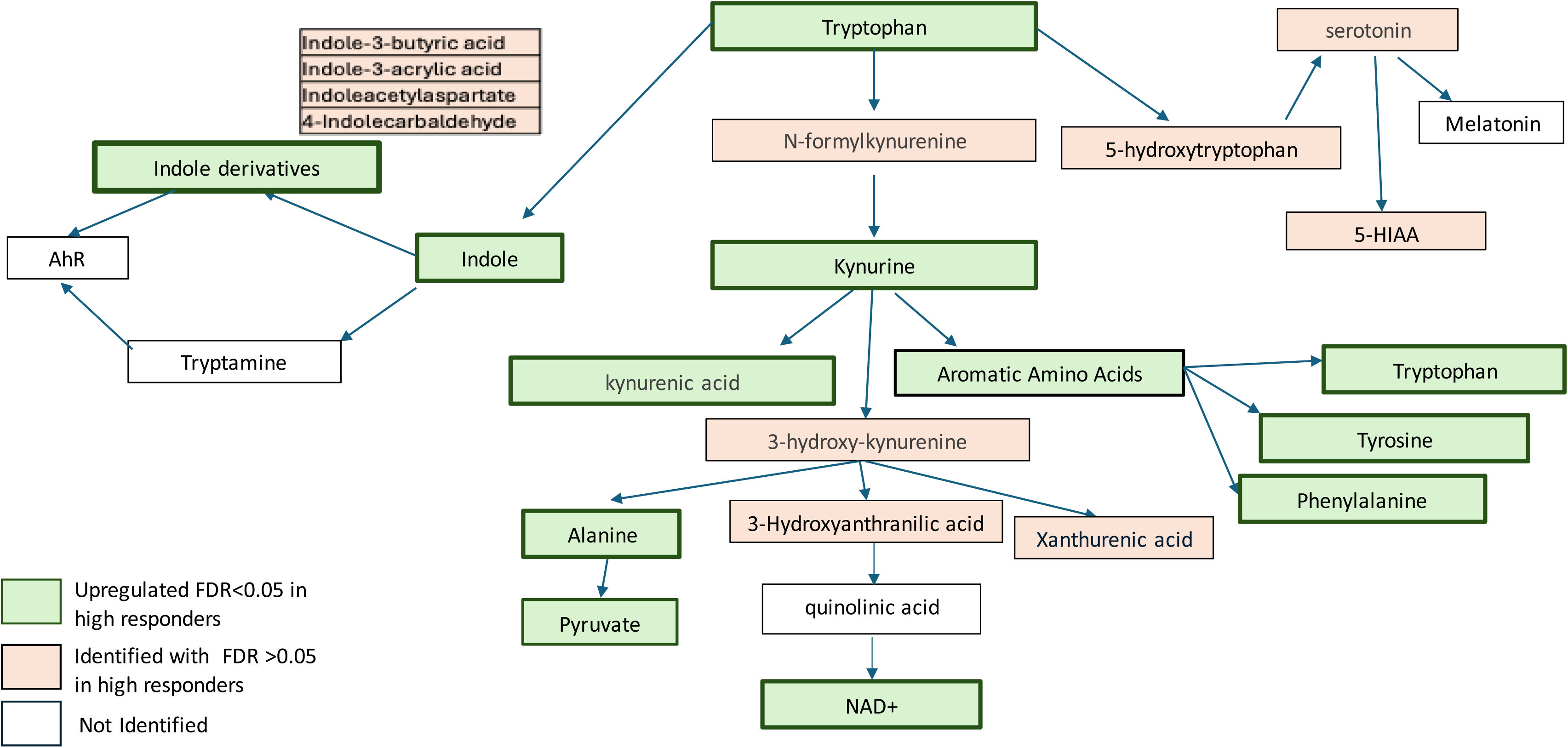
Tryptophan-related pathways from *a*dapted from the KEGG database. The FDRs were calculated using multiple t-tests in MetaboAnalyst 6.0. A large number of tryptophan metabolites were identified in muscle tissue and found to be upregulated in the high-responder group. At this stage, to further elaborate the tryptophan metabolism function, we extracted the KEGG IDs that were identified from Mummichog pathways and were introduced into Cytoscape_Metscape version 3.10.2. B) Tryptophan-kynurenine metabolism.

**Figure 7.**
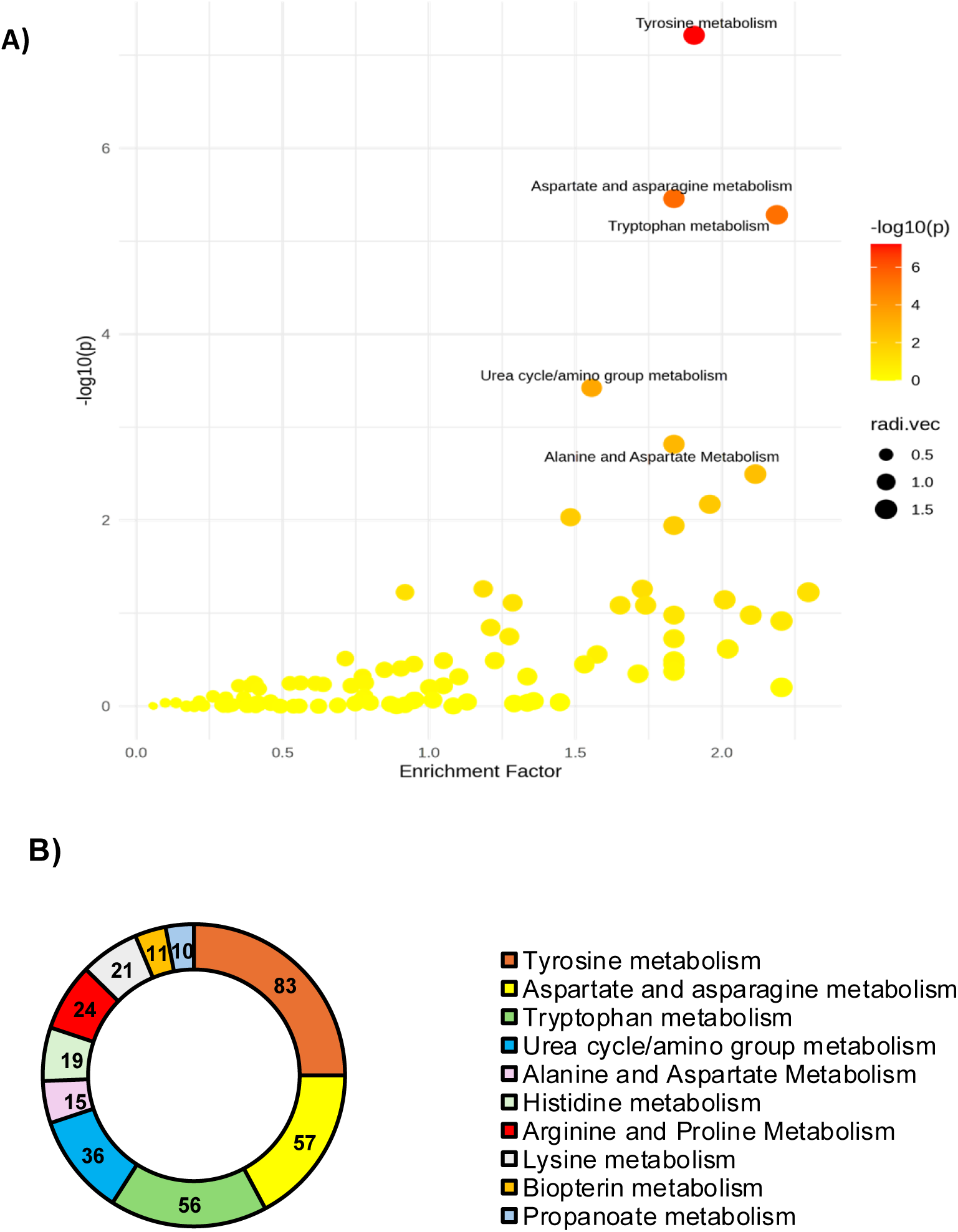
Mummichog analysis. A) Displays the results from the Mummichog analysis conducted using MetaboAnalyst 6.0. Unique identified m/z values with p-values set to the top 10% threshold (0.008) were used, along with retention time (in minutes) and analysis mode (positive), to generate the enriched pathways. The MFN human genome-scale metabolic model, which is manually curated and derived from multiple sources (KEGG, BIGG, Edinburgh Model, and Recon2), was utilized. The most enriched pathways identified include tyrosine metabolism, aspartate and arginine metabolism, tryptophan metabolism, the urea cycle, and amino acid groups. B) Shows the top 10 enriched metabolic pathways from Mummichog analysis with P-Fisher 0.05, and numbers in the ring represent the number of significant metabolite hits.

## 4 DISCUSSION

The findings of this study revealed that the metabolic profile of older individuals exhibited different responses to RT training, aligning with a phenotype characterized by a greater anabolic response (HighR). We propose that these findings offer important insight into the phenomenon of anabolic resistance, which is characterized by a poorer response to the normally robustly anabolic stimuli of RT and feeding [8]. Here, we provide a comprehensive interpretation of the significant metabolites that were identified and differed between individuals who showed a robust anabolic response versus a lesser anabolic response based on their known and potential functions in muscle metabolism, exercise and ageing.

### 4.1 Branched-chain amino acids

We showed that valine, leucine, and isoleucine were elevated in the skeletal muscle of HighR (Figure 3). This finding is in agreement with the notion that the expression of BCAA aminotransferase (BCAT2), an enzyme that catabolizes the first step of BCAA transamination, is highly expressed and active in the skeletal muscle [14]. We have previously reported and shown a key role for the BCAT2 gene in relation to skeletal muscle hypertrophy [14]. One factor contributing to this phenomenon with resistance exercise could be the upregulation of BCAT2, leading to the production of keto-acids of each BCAA [15]. This phenomenon may be reversed in resistance-trained individuals who often exhibit higher BCAA levels in their skeletal muscle, partly due to increased skeletal muscle insulin sensitivity. A recent study using mouse models validated that impaired BCAA catabolism weakens muscle mass and strength by disrupting mTOR signaling. Additionally, enhancing BCAA catabolism with BT2 was found to protect against sarcopenia in aged mice and in mice lacking Ppm1k, a key positive regulator of BCAA catabolism in skeletal muscle[16]. We also propose that HighR individuals may possess more efficient BCAA metabolism, facilitating the conversion of BCAAs into branched-chain keto acids (BCKAs), which may be more closely related to the regulation of muscle protein turnover (Figure 5). While knowledge regarding their role is limited, enhanced fatty acid oxidation has been shown to increase the ratio of mitochondrial acetyl-CoA:CoA and NADH:NAD+, leading to the inactivation of pyruvate dehydrogenase [17]. In contrast, there is limited knowledge regarding the functions of branched-chain α-ketoacid dehydrogenase (BCKDH) complex in human muscle with RT.

### 4.2 Tryptophan metabolism

Our pathway and functional analysis showed significant enrichment of tryptophan metabolism in HighR compared to LowR (Figure 6). The tryptophan pathway is involved in inflammation, immune responses, and excitatory neurotransmission; furthermore, it has been implicated in various diseases [18]. Tryptophan is an essential amino acid, which is degraded by kynurenine enzymes (e.g., indoleamine 2,3-dioxygenase and/or kynurenine aminotransferase) into kynurenines and is critical for the modulation of muscle protein synthesis, as well as the immune and central nervous system function [19]. The kynurenine pathway of tryptophan degradation is the major catabolic pathway for this essential amino acid [20]. Our results confirm previous work, where the muscle kynurenine levels were higher (Figure 4) in the muscle tissue of active vs. sedentary older adults [21]. At the same time, its downstream metabolites, kynurenic acid and nicotinamide adenine dinucleotide (NAD^+^) were also linked to better cardiorespiratory fitness and muscle oxidative capacity [18]. Further, tryptophan metabolism is also regulated in the gut, especially in the production of indole-derived metabolites [22-24]. Work in murine models and C2C12 cells suggested the role of indole propionic acid in protecting against inflammation [25], and the microbial indole has been shown to affect growth and metabolic function in multiple organs, including muscle, in mutated indole-producing wild-type mice [26]. Overall, our findings aligned with the previous models, suggesting that there is an important exercise–gut–muscle interaction regulated by tryptophan metabolism.

### 4.3 Gut-derived metabolites

In addition to tryptophan metabolism, we identified metabolites such as 4-hydroxyhippurate, creatine, proline, and stachydrine that were upregulated in the High R (Figure 3). Interestingly, recent Mendelian randomization analysis (utilizing gut-related metabolites genes) has shown a causal link between these metabolites with muscle function and muscle mass. Further, analysis using faecal samples has shown an association of several bacterial metabolites and increased lipopolysaccharide biosynthesis with depleted phenylalanine, tyrosine, and tryptophan biosynthesis in older adults with sarcopenia [27]. In addition, cholic acid was also upregulated, which is a bile acid utilized in the gut (Figure 4)[28]. Further interventional studies focusing on the gut-muscle axis are needed to substantiate these findings, especially on how RT relates to the upregulation/downregulation of gut-related metabolites may be warranted.

### 4.4 Carnosine

Our study showed a higher relative abundance of carnosine in HighR compared to LowR (Figureure 3C). In humans, carnitine is produced primarily in the liver and kidneys from the amino acids lysine and methionine, with the help of vitamins C, B6, and niacin [29]. Carnosine is a naturally occurring dipeptide composed of two amino acids, beta-alanine and histidine, and it is found in high concentrations in skeletal muscle [30]. Higher levels of muscle carnosine and carnitine are associated with improved muscle function, intracellular pH buffering capacity, ATP regeneration, and energy availability [31], which are critical for muscle adaptation following RT, explaining, in part, the greater capacity of HighR to gain muscle. Hoetker et al. [32] showed mitochondrial carnitine homeostasis (m-carn) contents and ATPDG1 gene expression concomitantly fluctuate throughout different phases of exercise, suggesting that the amount of carnosine synthesis is an important regulator of m-carn homeostasis. In general, our findings show that concomitant upregulation of carnosine synthesis is likely involved in maintaining stable carnitine levels after 10 weeks of RT, which could indicate a more efficient energy production and fatigue resistance in HighR compared to LowR.

### 4.5 Acylcarnitine

We observed that RT exercise affected muscle levels of acylcarnitines (a short-chain acylcarnitine [C2]) in the HighR compared to LowR. There are more than 1000 types of acylcarnitines in the human body. The general role of acylcarnitines is to transport acyl-groups (organic acids and fatty acids) from the cytoplasm into the mitochondria so that they can be broken down to produce energy [21, 33]. Our findings align with previous work suggesting that exercise can induce upregulation of acylcarnitines in humans, which influences muscle bioenergetics and acetyl group balance during and after exercise [34]. The key acylcarnitines identified in the current study are shown in Supplementary Table 3, include acetylcarnitine, a short-chain metabolite involved in energy metabolism, and 3-hydroxyoctanoylcarnitine, a medium-chain hydroxylated acylcarnitine indicative of incomplete fatty acid β-oxidation. The medium-chain acylcarnitines, including O-heptanoylcarnitine (C7) and carnitine (C8), play an important role in the carnitine shuttle [33].

### 4.6 Neurotransmitters

Denervated muscle fibres have been proposed as another contributor to declines in muscle mass and strength during ageing, for which altered acetylcholine (ACh) receptors have been implicated in partially explaining neuromuscular junction instability [5]. Eight weeks of heavy RT in healthy older men have previously shown a decrease in Ach receptor subunits α1 and ε subunit messenger RNA (mRNA), which were accompanied by an increase in muscle strength and (type II fibre) hypertrophy [5]. However, it is worth noting that receptor subunit levels do not directly reflect ACh production or release. The increased muscle ACh levels of HighR may indicate an enhanced ACr release at the neuromuscular junction of vastus lateralis (Figure 4). We have found several other neurotransmitters (Figure 4C), which confirm the important role of motor unit engagement in better response to RT in older people.

### 4.7 Pathways and functional analyses

Overall, we showed that muscle is an active metabolism organ where we identified more than 100 enriched pathways as a result of loading exercise. Overall, these pathways seem to have a significant role in the mechanisms of ageing, muscle weakness and frailty. Recently, Pu et al.[35] reported plasma NMR metabolomics and identified glycolytic and gluconeogenic metabolites, including lysine, leucine, and Acetoacetyl-CoA, to be related to frailty index, which has been identified in our muscle LC-MS analysis. Further, a large-scale study utilizing plasma mass-spectrometry data showed carnitine shuttle pathways [6], which was aligned with our study, where 11 significant hits of carnitine shuttle metabolites were upregulated in response to RT.

### 4.8 Considerations

Some important considerations must be acknowledged when interpreting our findings. The muscle samples were taken 48 hours after training, and thus, the metabolic shift predominantly reflected the medium to longer-term effect of exercise on muscle [36]. Dietary intake of participants was monitored before RT, and during RT, and there was no significant difference between the groups, which is a strength of the study design [8]. To fully uncover the effects of RT in older adults, further data on circulating metabolites and faecal microbiota could provide a more comprehensive picture of metabolomic changes. Given the sample size, we did not perform a sex-based analysis. Despite this, we had males and females in the LowR and HighR groups. A common question in ‘responder-based’ analysis is whether participants that are characterised as responders are persistently responders. In this case, a strategy might be to let all participants detrain and retrain them to assess this. Such an approach is highly impractical, time-consuming and inordinately expensive; however, we note that our classification into the top and bottom 25 responders with 4 sets of exercise was preserved in the contralateral limb that performed only 1 set of exercise in the same subjects. That is, the physiological hypertrophy response was conserved within an individual and so we propose our responder status was robust.

## 5 CONCLUSIONS

Our results contribute to understanding the mechanistic differences between HighR and LowR to RT in ageing. These data also provide an opportunity for future interventions that could guide the development of studies (i.e., dietary interventions) aiming to optimize muscle health and reduce the burden of sarcopenia and frailty in older adults. Additionally, the changes observed between groups may be used to monitor muscle adaptations to RT, which could help predict hypertrophic responses over time. This information could assist in adjusting their therapeutic strategies to support LowR individuals by initiating additional nutritional or nutraceutical interventions that may be beneficial.

## Supporting information

Mumichog Results

## Acknowledgments

This study was supported by the UKRI Biotechnology and Biological Sciences Research Council (MI, HM, SMP). SMP was supported by the Canada Research Chairs Program (CRC-2021-00495). The authors are grateful to Fundação de Amparo à Pesquisa do Estado de São Paulo (FAPESP)—Process Numbers: 2016/22635-6 and 2018/15691-2; and Conselho Nacional de Desenvolvimento Científico e Tecnológico (CNPq) for financial support. The authors would also like to thank the Participating Investigators for their contriubution in collecting the data, supporting the study particpants, techical editing of the raw data, and metabolomic data prcoessing.

## Data availability

The data that support the findings of this study are available from the corresponding author upon reasonable request. The data are not publicly accessible because of information that could compromise the privacy or consent of research participants.

## Author contributions

CL, ML, DT, YX, HR, KP, SMP, HM, MI have been involved in drafting the manuscript or revising it critically for important intellectual content. All authors have given final approval of the version to be published. All authors agreed to be accountable for all aspects of the work in ensuring that questions related to the accuracy or integrity of any part of the work are appropriately investigated and resolved. All authors have made substantial contributions to conception and design, and acquisition of data, analysis and interpretation of data. All authors have made substantial contributions to conception and design, and acquisition of data, analysis and interpretation of data.

## References

1. Thomas, A.C.Q., et al., Exercise-specific adaptations in human skeletal muscle: Molecular mechanisms of making muscles fit and mighty. Free Radical Biology and Medicine, 2024. 223: p. 341–356.

2. Krzysztofik, M., et al., Maximizing Muscle Hypertrophy: A Systematic Review of Advanced Resistance Training Techniques and Methods. Int J Environ Res Public Health, 2019. 16(24).

3. Baumert, P., et al., Skeletal muscle hypertrophy rewires glucose metabolism: An experimental investigation and systematic review. J Cachexia Sarcopenia Muscle, 2024. 15(3): p. 989–1002.

4. Gehlert, S., et al., Effects of Acute and Chronic Resistance Exercise on the Skeletal Muscle Metabolome. Metabolites, 2022. 12(5).

5. Soendenbroe, C., et al., Human skeletal muscle acetylcholine receptor gene expression in elderly males performing heavy resistance exercise. American Journal of Physiology-Cell Physiology, 2022. 323(1): p. C159–C169.

6. Rattray, N.J.W., et al., Metabolic dysregulation in vitamin E and carnitine shuttle energy mechanisms associate with human frailty. Nat Commun, 2019. 10(1): p. 5027.

7. Deelen, J., et al., A metabolic profile of all-cause mortality risk identified in an observational study of 44,168 individuals. Nat Commun, 2019. 10(1): p. 3346.

8. Lixandrão, M.E., et al., Higher resistance training volume offsets muscle hypertrophy nonresponsiveness in older individuals. J Appl Physiol (1985), 2024. 136(2): p. 421–429.

9. Broadhurst, D., et al., Guidelines and considerations for the use of system suitability and quality control samples in mass spectrometry assays applied in untargeted clinical metabolomic studies. Metabolomics, 2018. 14(6): p. 72.

10. Dunn, W.B., et al., The importance of experimental design and QC samples in large-scale and MS-driven untargeted metabolomic studies of humans. Bioanalysis, 2012. 4(18): p. 2249–64.

11. Mullard, G., et al., Erratum to: A new strategy for MS/MS data acquisition applying multiple data dependent experiments on Orbitrap mass spectrometers in non-targeted metabolomic applications. Metabolomics, 2015. 11(5): p. 1081–1081.

12. Xu, Y. and R. Goodacre, On Splitting Training and Validation Set: A Comparative Study of Cross-Validation, Bootstrap and Systematic Sampling for Estimating the Generalization Performance of Supervised Learning. J Anal Test, 2018. 2(3): p. 249–262.

13. Thévenot, E.A., et al., Analysis of the Human Adult Urinary Metabolome Variations with Age, Body Mass Index, and Gender by Implementing a Comprehensive Workflow for Univariate and OPLS Statistical Analyses. J Proteome Res, 2015. 14(8): p. 3322–35.

14. Mann, G., et al., Branched-chain amino acids: catabolism in skeletal muscle and implications for muscle and whole-body metabolism. Frontiers in Physiology, 2021. 12: p. 702826.

15. Chaillou, T., et al., Time course of gene expression during mouse skeletal muscle hypertrophy. Journal of applied physiology, 2013. 115(7): p. 1065–1074.

16. Zuo, X., et al., Multi-omic profiling of sarcopenia identifies disrupted branched-chain amino acid catabolism as a causal mechanism and therapeutic target. Nat Aging, 2025.

17. Savage, D.B., K.F. Petersen, and G.I. Shulman, Disordered lipid metabolism and the pathogenesis of insulin resistance. Physiol Rev, 2007. 87(2): p. 507–20.

18. Xue, C., et al., Tryptophan metabolism in health and disease. Cell Metab, 2023. 35(8): p. 1304–1326.

19. Martin, K.S., M. Azzolini, and J.L. Ruas, The kynurenine connection: How exercise shifts muscle tryptophan metabolism and affects energy homeostasis, the immune system, and the brain. American Journal of Physiology-Cell Physiology, 2020.

20. Agudelo, L.Z., et al., Kynurenic Acid and Gpr35 Regulate Adipose Tissue Energy Homeostasis and Inflammation. Cell Metab, 2018. 27(2): p. 378-392.e5.

21. Hinkley, J.M., et al., Exercise and ageing impact the kynurenine/tryptophan pathway and acylcarnitine metabolite pools in skeletal muscle of older adults. J Physiol, 2023. 601(11): p. 2165–2188.

22. Wang, Y.-C., et al., Indole-3-Propionic Acid Protects Against Heart Failure With Preserved Ejection Fraction. Circulation Research, 2024. 134(4): p. 371–389.

23. Enoki, Y., et al., Indoxyl sulfate potentiates skeletal muscle atrophy by inducing the oxidative stress-mediated expression of myostatin and atrogin-1. Sci Rep, 2016. 6: p. 32084.

24. Serger, E., et al., The gut metabolite indole-3 propionate promotes nerve regeneration and repair. Nature, 2022. 607(7919): p. 585–592.

25. Du, L., et al., Indole-3-Propionic Acid, a Functional Metabolite of Clostridium sporogenes, Promotes Muscle Tissue Development and Reduces Muscle Cell Inflammation. Int J Mol Sci, 2021. 22(22).

26. Xing, P.Y., et al., Microbial Indoles: Key Regulators of Organ Growth and Metabolic Function. Microorganisms, 2024. 12(4).

27. Wang, Y., et al., Population-based metagenomics analysis reveals altered gut microbiome in sarcopenia: data from the Xiangya Sarcopenia Study. Journal of cachexia, sarcopenia and muscle, 2022. 13(5): p. 2340–2351.

28. Aliwa, B., et al., Altered gut microbiome, bile acid composition and metabolome in sarcopenia in liver cirrhosis. J Cachexia Sarcopenia Muscle, 2023. 14(6): p. 2676–2691.

29. Vaz, F.M. and R.J. Wanders, Carnitine biosynthesis in mammals. Biochem J, 2002. 361(Pt 3): p. 417–29.

30. Boldyrev, A.A., G. Aldini, and W. Derave, Physiology and pathophysiology of carnosine. Physiol Rev, 2013. 93(4): p. 1803–45.

31. Li, G., Z. Li, and J. Liu, Amino acids regulating skeletal muscle metabolism: Mechanisms of action, physical training dosage recommendations and adverse effects. Nutrition & Metabolism, 2024. 21(1): p. 41.

32. Hoetker, D., et al., Exercise alters and β-alanine combined with exercise augments histidyl dipeptide levels and scavenges lipid peroxidation products in human skeletal muscle. J Appl Physiol (1985), 2018. 125(6): p. 1767–1778.

33. Dambrova, M., et al., Acylcarnitines: Nomenclature, Biomarkers, Therapeutic Potential, Drug Targets, and Clinical Trials. Pharmacol Rev, 2022. 74(3): p. 506–551.

34. Seiler, S.E., et al., Carnitine Acetyltransferase Mitigates Metabolic Inertia and Muscle Fatigue during Exercise. Cell Metab, 2015. 22(1): p. 65–76.

35. Pu, Y., et al., Gut microbial features and circulating metabolomic signatures of frailty in older adults. Nat Aging, 2024.

36. Schranner, D., et al., Physiological extremes of the human blood metabolome: A metabolomics analysis of highly glycolytic, oxidative, and anabolic athletes. Physiol Rep, 2021. 9(12): p. e14885.

