## Supplementary Table 1 for "Global skeletal muscle metabolomics reveals mechanisms behind higher response to resistance training in older adults"

| **Supplementary Table 1. Orbitrap ID-X Tribid MS parameters** | | | | |
| --- | --- | --- | --- | --- |
| MS parameter | Polar (ESI+) | Polar (ESI-) | Nonpolar (ESI+) | Nonpolar (ESI-) |
| m/z range | 66.7 - 1000 | 66.7 - 1000 | 150-2000 | 150-2000 |
| Resolution (FWHM at m/z 200) | 120,000 | | 60,000 | |
| Spray voltage (kV) | 3.2 | 3.0 | 3.2 | 2.7 |
| Sheath Gas (AU) | 40 | | 40 | |
| Aux Gas (AU) | 8 | | 8 | |
| Sweep Gas (AU) | 1 | | 1 | |
| Capillary Temp (℃) | 275 | | 300 | |
| ESI Heater Temp (℃) | 320 | | 350 | |
| Max. Injection time (s) | 100 | | 50 | |
