## Supplementary Table 2 for "Global skeletal muscle metabolomics reveals mechanisms behind higher response to resistance training in older adults"

| **Supplementary Table 2. UHPLC conditions** | | | | |
| --- | --- | --- | --- | --- |
| UHPLC conditions | Polar | | Non-polar | |
| Flow rate | 0.4mL/min | | | |
| Buffer composition | 0.1% Formic Acid (FA) | | 10mM Ammonium Formate and 0.1% Formic Acid (FA) | |
| Solvent composition | %(A) | %(B) | %(A) | %(B) |
| Time (mins) | Gradient curve = 5, linear | | | |
| 0 | 80% | 20% | 80% | 20% |
| 1.60 | 80% | 20% | 80% | 20% |
| 9.40 | 100% | 0% | 100% | 0% |
| 10.60 | 100% | 0% | 100% | 0% |
| 12.60 | 80% | 20% | 80% | 20% |
| 15.00 | 80% | 20% | 80% | 20% |
| 15.00 | Gradient end | | | |
