## Supplementary Table 3 for "Global skeletal muscle metabolomics reveals mechanisms behind higher response to resistance training in older adults"

| Supplementary Table 3. Acylcarnitine metabolites | | | | | | | | |  |  |
| --- | --- | --- | --- | --- | --- | --- | --- | --- | --- | --- |
| Name | Formula | Annot. DeltaMass [Da] | Annot. DeltaMass [ppm] | Calc. MW | m/z | RT [min] | p-values | FDR (before and after | p-values | FDR |
| 3-Hydroxyhexadecadienoylcarnitine | C10 H19 N O5 | 0.00079 | 1.91 | 411.29926 | 412.30654 | 9.644 | 0.10890483 | 0.743574374 | 0.18770527 | 0.26797236 |
| [Similar to: Acetyl-L-carnitine; ΔMass: -0.9831 Da] | C11 H21 N O5 |  |  | 202.13268 | 203.13996 | 0.966 | 0.11538763 | 0.744888709 | 0.42749742 | 0.51585531 |
| 3-hydroxyoctanoylcarnitine | C13 H23 N O4 | 0.00055 | 1.8 | 303.20512 | 304.21239 | 6.44 | 0.13592501 | 0.769495728 | 0.03255324 | 0.07058424 |
| 3-Hydroxy-cis-5-tetradecenoylcarnitine | C15 H29 N O4 | 0.00072 | 1.87 | 385.28354 | 386.29082 | 8.57 | 0.50179443 | 0.912662865 | 0.01998473 | 0.04966156 |
| 3-hydroxytetradecanoylcarnitine | C15 H29 N O5 | 0.00083 | 2.14 | 387.2993 | 388.30658 | 8.958 | 0.50185487 | 0.912662865 | 0.00579639 | 0.02087634 |
| 3-Methylglutarylcarnitine | C17 H31 N O4 | 0.00039 | 1.34 | 289.15292 | 290.1602 | 3.872 | 0.54100513 | 0.919934016 | 0.01820953 | 0.04643358 |
| trans-2-Dodecenoylcarnitine | C17 H33 N O4 | 0.00063 | 1.85 | 341.25724 | 342.26452 | 9.3 | 0.60101875 | 0.931985646 | 0.08734244 | 0.14894497 |
| 2-Hexenoylcarnitine | C19 H35 N O4 | 0.00043 | 1.68 | 257.16314 | 258.17042 | 4.62 | 0.62941622 | 0.939878251 | 0.00536305 | 0.01971681 |
| trans-2-Dodecenoylcarnitine | C19 H35 N O4 | 0.00065 | 1.9 | 341.25726 | 342.26453 | 7.067 | 0.61716986 | 0.937667606 | 0.0318004 | 0.06927376 |
| 3-hydroxydodecanoylcarnitine | C19 H37 N O4 | 0.00066 | 1.82 | 359.26783 | 360.2751 | 9.07 | 0.67623259 | 0.952459655 | 0.04903524 | 0.09603707 |
| O-heptanoylcarnitine | C19 H37 N O5 | 0.00049 | 1.81 | 273.1945 | 296.18374 | 8.09 | 0.68649067 | 0.954590141 | 0.01961506 | 0.04898515 |
| 3-Methylglutarylcarnitine | C21 H37 N O5 | 0.00054 | 1.88 | 289.15308 | 290.16036 | 4.552 | 0.754735 | 0.964610089 | 0.00579668 | 0.02087634 |
| trans-2-Dodecenoylcarnitine | C21 H39 N O4 | 0.0006 | 1.76 | 341.25721 | 342.26449 | 9.379 | 0.7559905 | 0.964610089 | 0.07050005 | 0.1264279 |
| Acetylcarnitine | C21 H39 N O5 | 0.0008 | 2.05 | 387.29927 | 388.30655 | 8.738 | 0.82420314 | 0.975357909 | 0.00041864 | 0.00404212 |
| Carnitine (C8) | C21 H41 N O5 | 0.00025 | 0.88 | 287.20991 | 288.21719 | 7.676 | 0.87331381 | 0.978416351 | 0.75893662 | 0.81400968 |
| O-heptanoylcarnitine | C23 H41 N O5 | 0.00048 | 1.77 | 273.19449 | 274.20175 | 8.163 | 0.87188548 | 0.978416351 | 9.07E-05 | 0.0015904 |
